# *PRX9* and *PRX40* are extensin peroxidases essential for maintaining tapetum and microspore cell wall integrity during *Arabidopsis* anther development: Extensin peroxidases prevent tapetum hypertrophy

**DOI:** 10.1101/319020

**Authors:** Joseph R. Jacobowitz, Jing-Ke Weng

**Affiliations:** Whitehead Institute for Biomedical Research, Cambridge, Massachusetts, 02142, United States; Department of Biology, Massachusetts Institute of Technology, Cambridge, Massachusetts, 02139, United States

## Abstract

Pollen and microspore development is an essential step in the life cycle of all land plants that generate male gametes. Within flowering plants, pollen development occurs inside of the anther. Here, we report the identification of two class III peroxidase-encoding genes, *PRX9* and *PRX40*, that are genetically redundant and essential for proper anther and pollen development in *Arabidopsis thaliana*. *Arabidopsis* double mutants devoid of functional *PRX9* and *PRX40* are male-sterile. The mutant anthers display swollen, hypertrophic tapetal cells and pollen grains, suggesting disrupted cell wall integrity. These phenotypes ultimately lead to nearly 100%-penetrant pollen degeneration upon anther maturation. Using immunochemical and biochemical approaches, we show that *PRX9* and *PRX40* are likely extensin peroxidases that contribute to the establishment of tapetal cell wall integrity during anther development. This work identifies *PRX9* and *PRX40* as the first extensin peroxidases to be described in *Arabidopsis* and highlights the importance of extensin cross-linking during plant development.

## Introduction

Pollen performs a critical step in the flowering plant life cycle by transporting the male gamete to the female ovule for fertilization. To do this, pollen must survive the journey from its origin in the anther to its destination atop the carpel. Wind-borne or insect-dispersed pollen may need to travel great distances to reach a suitable carpel. During its journey, pollen must endure a wide range of abiotic stresses, including UV radiation, extreme temperatures, and desiccation. To survive these challenges, pollen grains are protected by a unique and specialized cell wall.

The synthesis of the pollen wall was one of the most important innovations that enabled plant life on land. The pollen wall is highly structured and indispensable for pollen vitality. It is composed of two parts: an outer exine and an inner intine (Heslop-Harrison, 1968). The exine is composed of a uniquely robust polymer called sporopollenin. Many genes involved in the biosynthesis of sporopollenin have been identified, mostly through genetic studies, but the detailed chemical structure of the polymer is still poorly understood (Quilichini et al., 2015). The intine resembles a conventional plant cell wall and consists of cellulose, hemicellulose and pectin, as well as cell-wall-associated proteins (Knox et al., 1970). Pollen development occurs in the anther, concurrent with anther development.

*Arabidopsis* anthers have four functionally equivalent lobes that are designed to produce and release pollen grains. Within each lobe, pollen develops in a chamber known as the locule. Locule walls are lined by a specialized tissue composed of secretory cells known as the tapetum. The tapetum is the innermost layer of the anther and provides nutrients to developing pollen grains. The tapetum is also thought to supply the sporopollenin precursors that later polymerize to form the pollen exine. At late stages of pollen development, the tapetum undergoes programmed cell death, depositing a mixture of protein and wax on the surface of the exine, known as tryphine (Heslop-Harrison, 1968).

Although the sequence of events in tapetum and pollen development has been well-described, the exact biochemical processes involved remain poorly understood. In the model plant *Arabidopsis thaliana*, hundreds of evolutionarily conserved genes are specifically expressed in developing anthers, but only a few have been functionally characterized to date (Li et al., 2017). In this work, we sought to better understand the role of a subset of these genes encoding class III peroxidases.

Class III peroxidases are a large family of heme-iron-dependent peroxidases that arose specifically within land plants and has expanded widely since their emergence; the *Arabidopsis thaliana* genome contains 73 members (Tognolli et al., 2002; Duroux and Welinder, 2003; Welinder et al., 2002; Weng and Chapple, 2010). Many biochemical activities important for various aspects of land plant physiology have been attributed to class III peroxidases. For example, class III peroxidases oxidatively polymerize monolignols in the apoplast of the lignifying cells. Previous studies have found that *PRX2*, *PRX4*, *PRX17*, *PRX25*, *PRX52*, *PRX71*, and *PRX72* are involved in stem lignification in *Arabidopsis* (Shigeto et al., 2015; Fernández-Pérez et al., 2015b; Herrero et al., 2013; Fernández-Pérez et al., 2015a). In addition, class III peroxidases are also thought to polymerize other elements of the plant cell wall, including suberin and extensins (Bernards et al., 1999; Jackson et al., 2001). Conversely, some class III peroxidases may function to destabilize plant cell walls by generating hydroxyl radicals that cleave load-bearing cell wall polysaccharides (Liszkay et al., 2004). This is exemplified by *PER36/PRX36*, which serves as a seed mucilage extrusion factor by loosening the walls of epidermal cells in the *Arabidopsis* seed coat (Kunieda et al., 2013).

In this work, we identified two class III peroxidase-encoding genes, *PRX9* and *PRX40*, that are required for proper development of tapetum and microspores in *Arabidopsis*. The *prx9 prx40* double mutants exhibit distinctive tapetum swelling and enlarged developing pollen grains, ultimately leading to microspore degeneration and male sterility. We provide evidence to indicate that *PRX9* and *PRX40* are extensin peroxidases that ensure proper establishment of the tapetum and microspore cell walls.

## Results

### *PRX9* and *PRX40* are specifically expressed in the tapetum of developing anthers

To identify uncharacterized metabolic enzymes involved in pollen and anther development, we performed a co-expression analysis using the known exine biosynthetic genes as search queries (Obayashi et al., 2007). This analysis uncovered two class III peroxidases, *PRX9* (*AT1G44970*) and *PRX40* (*AT4G16270*), which are specifically expressed in early stages of flower development **(Figure S1)** (Waese and Provart, 2017; Winter et al., 2007).

To experimentally examine the tissue specificity of *PRX9* and *PRX40* expression, we first profiled the transcript abundance of *PRX9* and *PRX40* across six different *Arabidopsis* tissue types by RT-PCR. This experiment verified that *PRX9* and *PRX40* are both specifically expressed in the flower buds but not in the other five tissues examined (**Figure 1A**). To obtain higher spatial and temporal resolution for *PRX9* and *PRX40* expression, we generated promoter-β-Glucuronidase (GUS) reporter lines for *PRX9* and *PRX40* and examined the tissue-specific GUS activities in six independent T1 transformants for each reporter construct. In both cases, all six replicates produced nearly identical results. Whereas *PRX9* promoter-reporter activity was detected in both early developing anthers and mature carpels, *PRX40* promoter-reporter activity was exclusively observed in early developing anthers (**Figure 1B-D**). Closer examination of the GUS-stained anther sections further established that *PRX9 and PRX40* are specifically expressed in the tapetum (**Figure 1E-F**).

**Figure 1.**
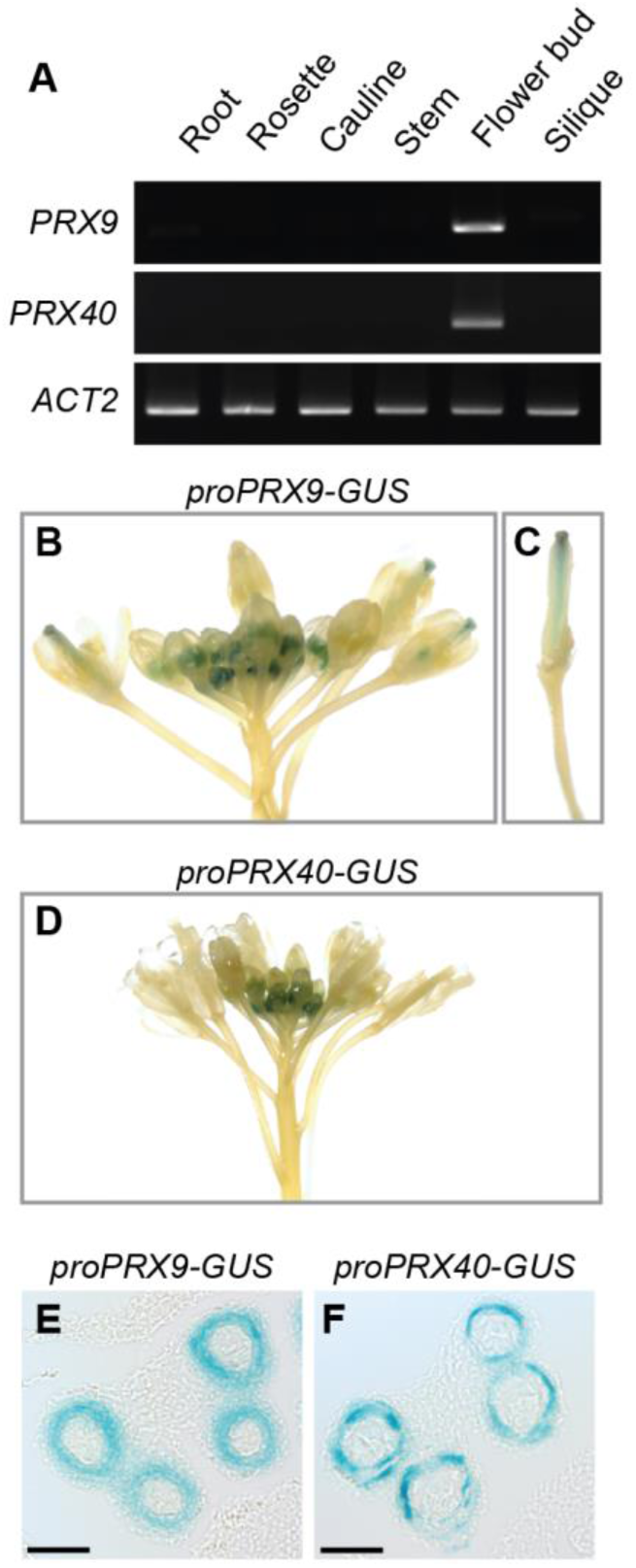
*PRX9* and *PRX40* are expressed in the tapetum of developing anthers. (**A**) RT-PCR across a variety of Col-0 tissues shows *PRX9* and *PRX40* are specifically expressed in the flower buds. Whole mount images show that reporter *proPRX9-GUS* activity is detected in the early anthers (**B**) and mature carpel (**C**). *proPRX40-GUS* reporter activity is detected in the early anthers (**D**). Images of anther sections showing reporter expression is tapetum-specific for both *proPRX9-GUS* (**E**) and *proPRX40-GUS* (**F**). Bars = 25 μm.

### *PRX9 and PRX40* are genetically redundant and required for male-fertility

To probe the *in vivo* function of *PRX9 and PRX40*, we obtained two independent T-DNA insertion lines for each of the two genes. SALK_204557 (*prx9-1*) and SAIL_875_A08 (*prx9-2*) contain insertions in the second and fourth exons of *PRX9*, respectively, and SALK_031680 (*prx40-*1) and SALK_061270 (*prx40-2*) contain insertions in the first and second introns of *PRX40*, respectively **(Figure 2A)**. To assess whether these insertion events resulted in loss-of-function mutations, we examined the expression of *PRX9* and *PRX40* in the T-DNA lines by RT-PCR (**Figure 2B)** and established that all the four T-DNA insertion events produced loss-of-function alleles.

**Figure 2.**
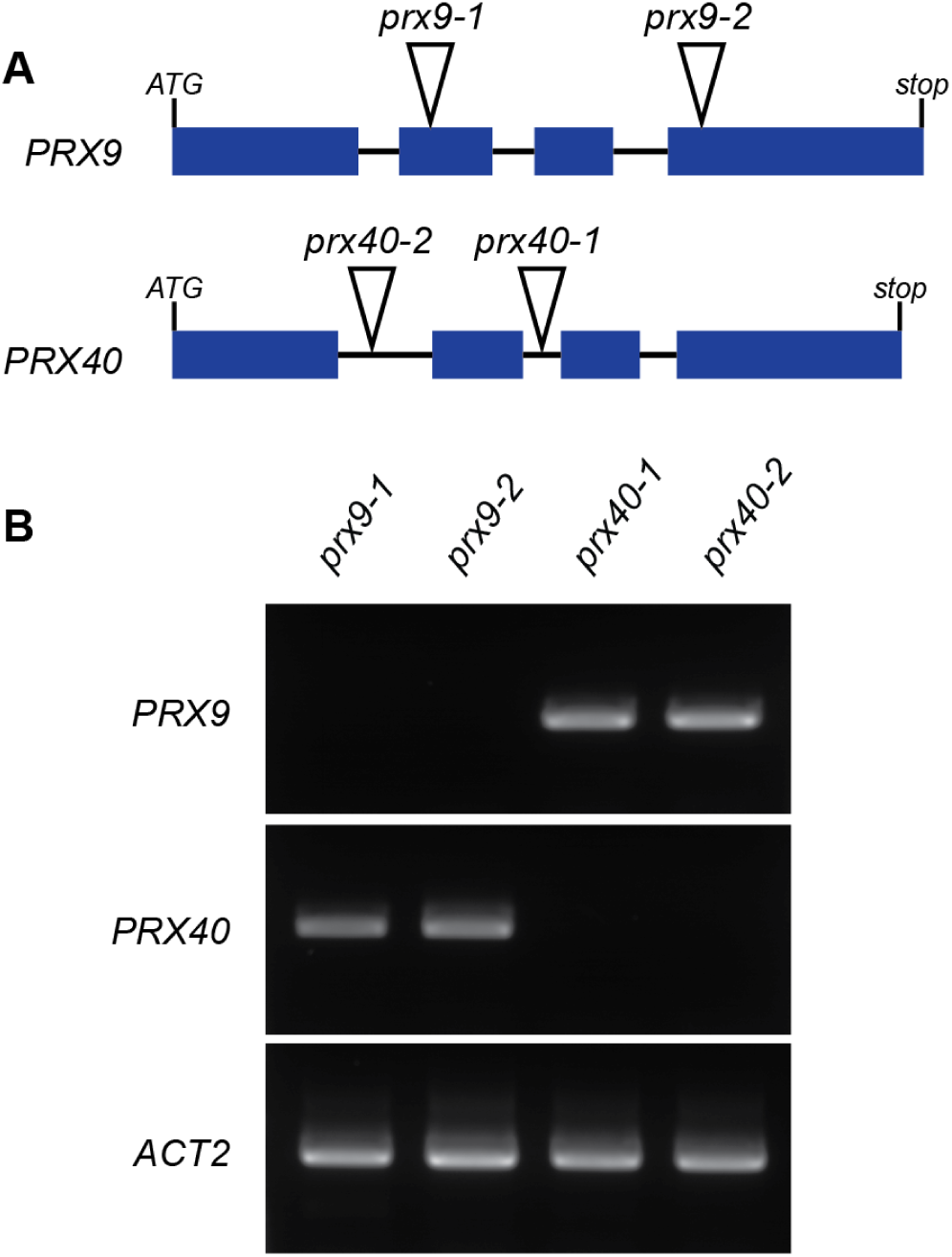
Molecular characterization of *PRX9* and *PRX40* insertion lines. (**A**) *prx9-1* and *prx9-2* contain T-DNA insertions in the second and fourth exons of *PRX9*, respectively. *prx40-1* and *prx40-2* contain T-DNA insertions in the second and first introns of *PRX40*, respectively. (**B**) RT-PCR showing full-length *PRX9* cDNA could not be amplified from *prx9-1* or *prx9-2* mutants, while full-length *PRX40* cDNA could not be amplified from *prx40-1* or *prx40-2 mutants*.

Single mutants of either *prx9* or *prx40* display no visible differences from Col-0 wild-type plants. In particular, the *prx9* and *prx40* mutants are fully fertile, and develop fully elongated, seed-bearing siliques (**Figure 3B-E**).

**Figure 3.**
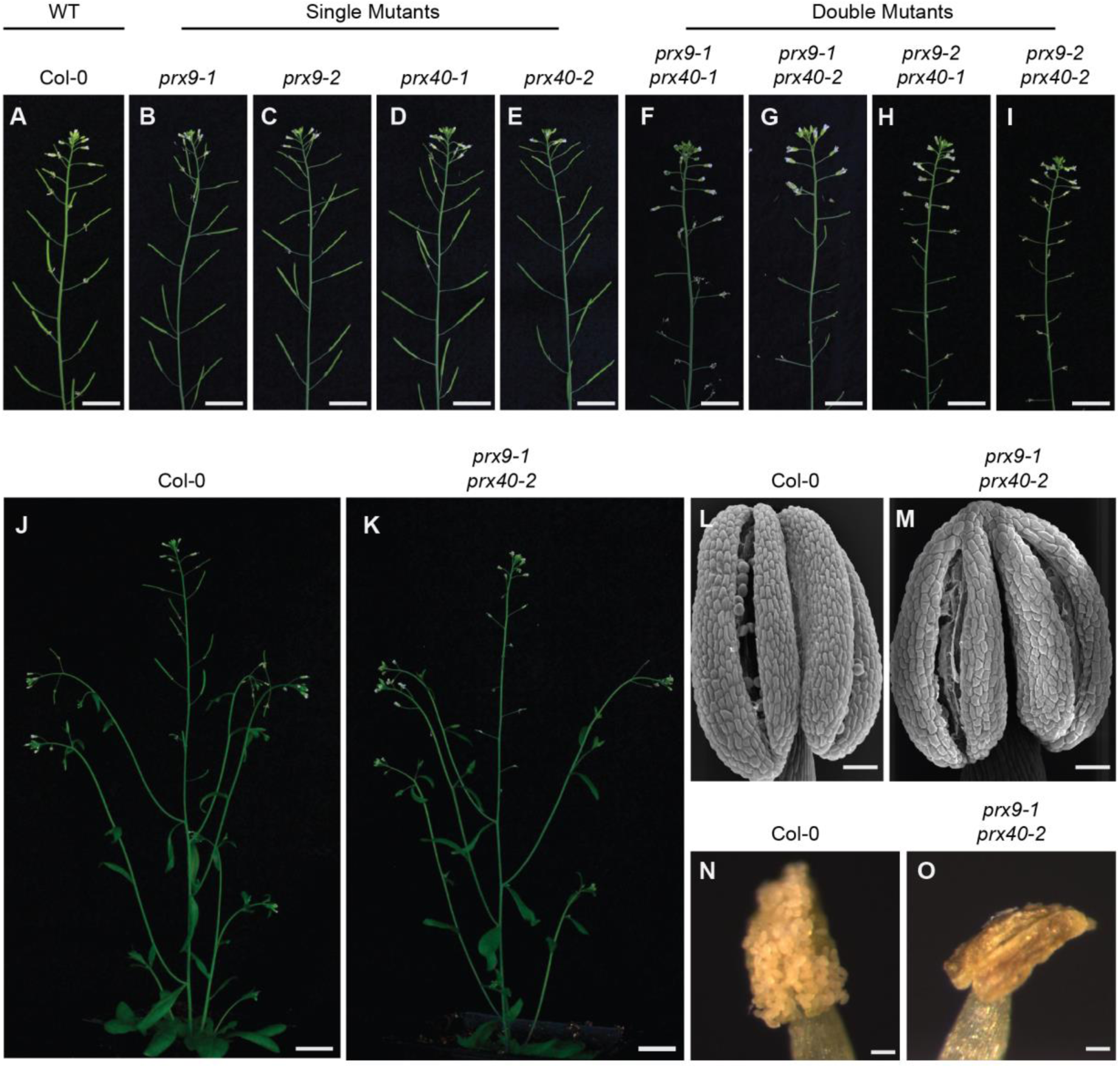
The *prx9 prx40* double mutants are male-sterile and do not produce viable pollen. While wild-type (**A, J**), single *prx9* (**B, C**), and single *prx40* (**D, E**) plants show no defects in fertility, all *prx9 prx40* double mutants (**F-I, K**) are male-sterile. Bars = 2 cm. Stage 13 wild-type (**L**) and *prx9-1 prx40-2* (**M**) anthers dehisce normally. Bars = 50 μm. Stage 14 wild-type (**N**) anthers are coated in pollen grains. *prx9-1 prx40-2* (**O**) anthers are devoid of pollen grains. Stages according to (Sanders et al., 1999). Bars = 50 μm.

Since none of the single mutants exhibited any visible phenotype, we hypothesized that *PRX9* and *PRX40* may be genetically redundant. To test this, we generated all four possible *prx9 prx40* double mutants using the four single mutant alleles. In contrast to the single mutants, which displayed no obvious defects in fertility, all *prx9 prx40* double mutant plants were completely sterile (**Figure 3 F-I**). Siliques from double mutants were short and contained no seeds. The *prx9 prx40* plants are likely male-sterile as *prx9 prx40* plants were readily fertilized by wild-type pollen, but all attempts to fertilize wild-type plants with *prx9 prx40* pollen failed. Since two independent alleles are represented for each peroxidase gene, and all four double mutants displayed identical phenotypes, we conclude that *PRX9* and *PRX40* are genetically redundant and required for male fertility in *Arabidopsis*.

We then examined the flowers from the double mutant at various developmental stages by light microscopy. At stage 14 of anther development (Sanders et al., 1999), wild-type anthers were coated with mature pollen grains, whereas *prx9-1 prx40-2* anthers were devoid of pollen **(Figure 3N and O)**. We also observed that anthers from the *prx9-1 prx40-2* plants displayed defects in anther filament elongation (**Figure S2**). However, this did not appear to be the primary cause of sterility as *prx9 prx40* anthers were incapable of depositing pollen on the stigma, even when manually crossed. Since indehiscence is one of the mechanisms that can lead to male-sterility even when pollen grains are properly developed (Sanders et al., 1999; Hao et al., 2014; Yang et al., 2017), we examined stage 13 anthers (Sanders et al., 1999; Hao et al., 2014; Yang et al., 2017) by scanning electron microscopy (SEM). Anthers from both wild-type and *prx9-1 prx40-2* plants clearly showed breakage along the stomiums, indicating that the male-sterility phenotype of *prx9-1 prx40-2* is unlikely to be caused by indehiscence **(Figure 3L and M)**.

### *PRX9* and *PRX40* play important roles in maintaining tapetum and micropore cell wall integrity

To identify the stage at which *prx9-1 prx40-2* anther and pollen development first derails, we performed a histological comparison between *prx9-1 prx40-2* and wild-type anthers over various developmental stages (**Figure 4**). At stage 5/6, *prx9-1 prx40-2* and wild-type anthers were indistinguishable. Noticeable differences first appeared at stages 8/9 with the double mutant exhibiting ectopically swollen tapetal cells that invade into the locular space. The swollen tapetum persisted through stage 10 in *prx9-1 prx40-2* plants. By stages 11 and 12, the tapetum had degenerated in both plants, but the pollen grains were shriveled and crushed in the *prx9-1 prx40-2* anthers. These observations suggest that the male-sterility of *prx9-1 prx40-2* plants is likely caused by tapetum swelling (i.e. tapetum hypertrophy) and subsequent pollen degeneration.

**Figure 4.**
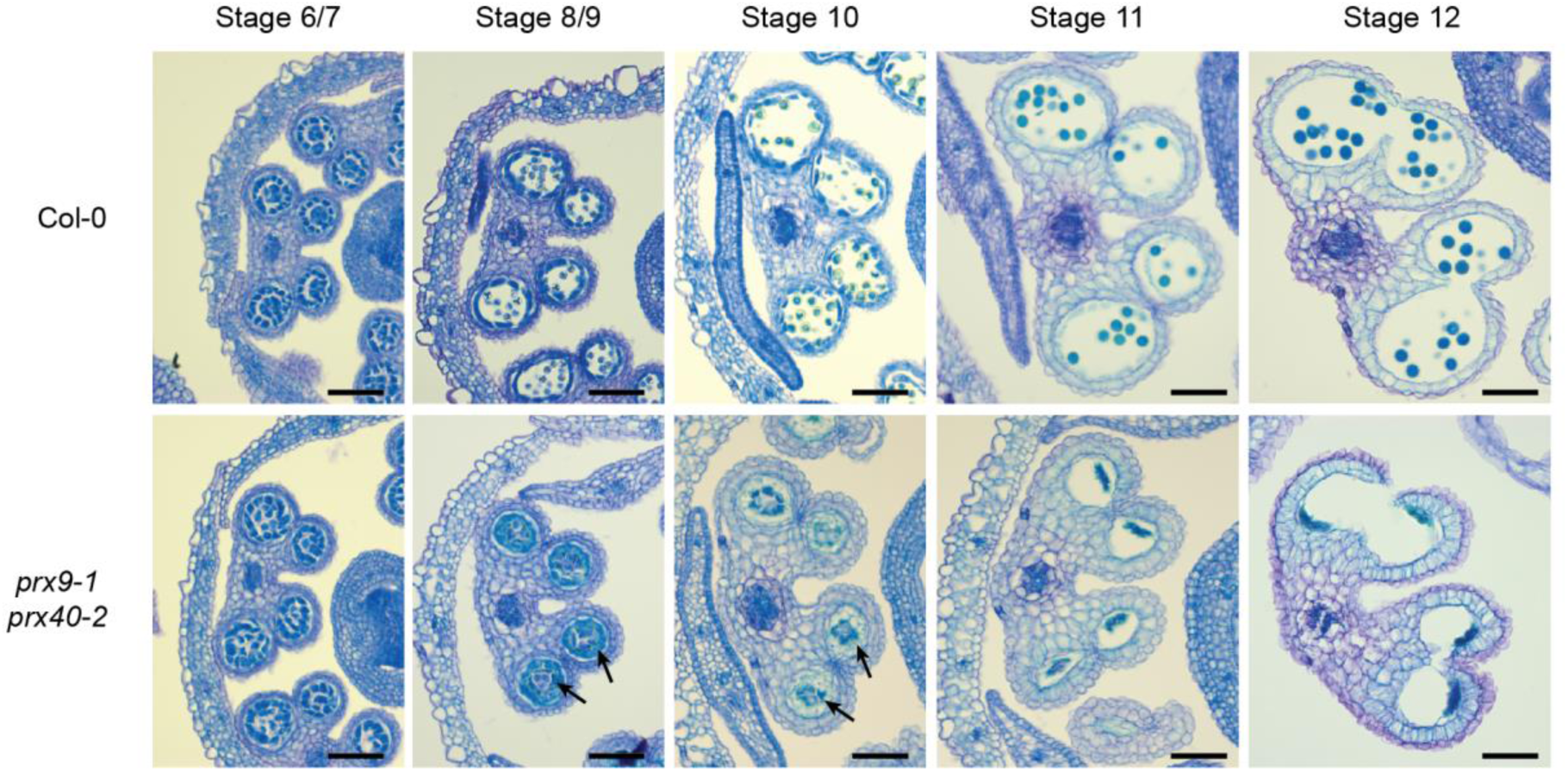
*prx9 prx40* anthers display tapetum hypertrophy and pollen degeneration. Wild-type (**Top Row**) and *prx9-1 prx40-2* (**Bottom Row**) anther development from stages 5 - 12 (Sanders et al., 1999). Tapetum swelling in *prx9-1 prx40-2* first appears at stage 8 and persists through stage 10, indicated by arrows. Tapetum programmed cell death and anther dehiscence occurs normally in both wild-type and *prx9-1 prx40-2* during stages 11 and 12. Bars = 25 μm.

To further characterize the tapetum hypertrophy phenotype, we examined stage 9 anthers by transmission electron microscopy (TEM) (**Figure 5**). Consistently, the *prx9-1 prx40-2* sample displayed expanded tapetum cells with enlarged tapetal vacuoles, when compared to wild type. Microspores in *prx9-1 prx40-2* anthers were also abnormally enlarged and vacuolated. The boundaries between tapetal cells in *prx9-1 prx40-2* plants were jigsaw-shaped, with one cell swelling into the neighboring cell, whereas the boundaries between tapetal cells in wild-type plants were typically well-defined and straight. This phenotype may be a manifestation of compromised tapetal cell wall integrity. Moreover, we also observed aberrant distributions of electron-dense material in the swollen tapetum of *prx9-1 prx40-2* anthers, reminiscent of sporopollenin typically observed at the outer wall of normally developed pollen. By stage 9, wild-type microspores exhibit a fully developed exine. By contrast, only nexine and probaculae had formed on the microspore surface in the double mutant with the majority of the sporopollenin-like material accumulated at the locular face of the tapetum. Finally, the middle layer was not crushed in *prx9-1 prx40-2* plants. Taken together, this suggests that *PRX9* and *PRX40* play an important role in maintaining tapetal and microspore cell size and shape by modulating the cell wall.

**Figure 5.**
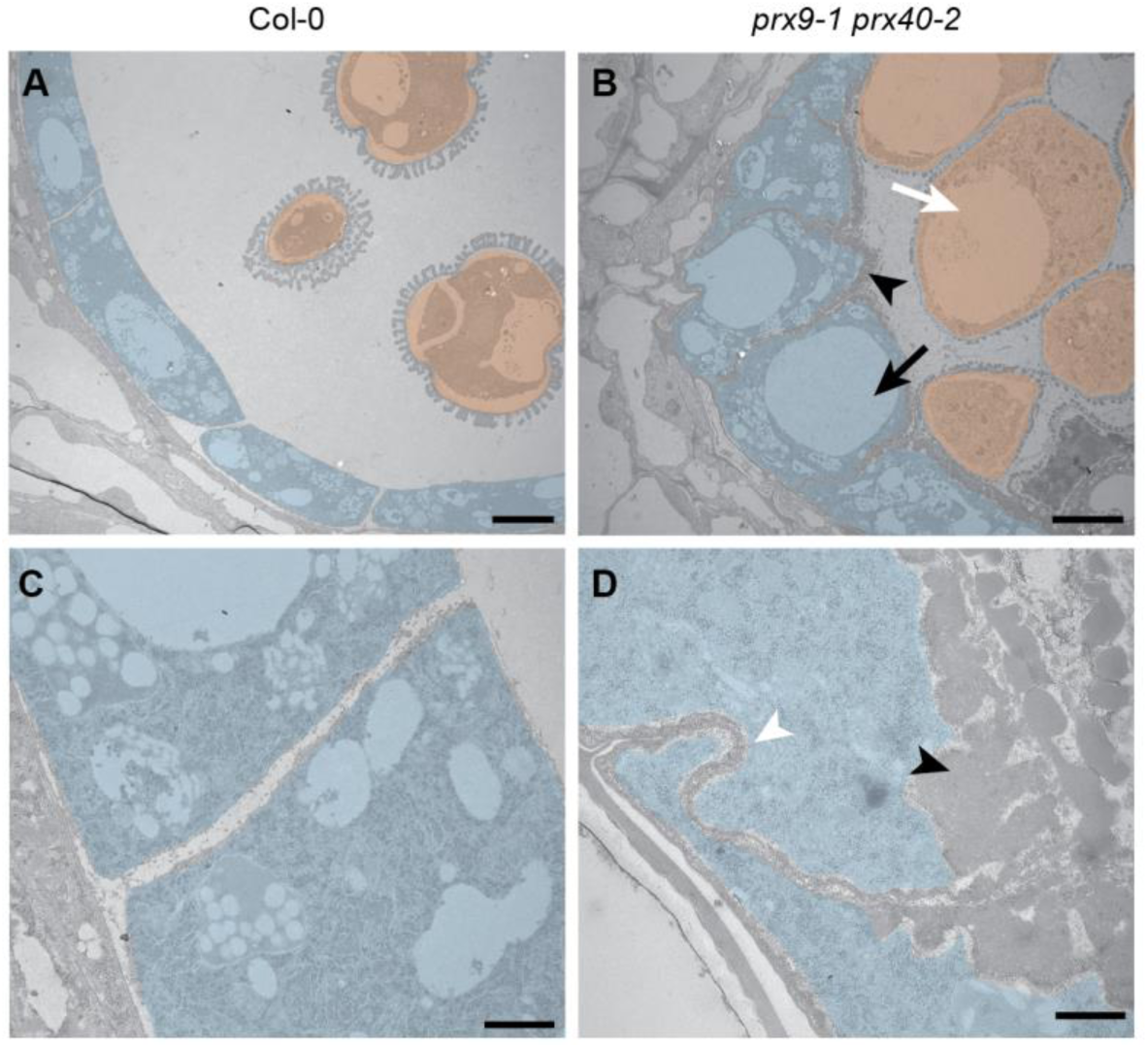
Stage 9 *prx9-1 prx40-2* anthers contain swollen, hyper-vacuolated tapetal cells and pollen grains. Stage 9 wild-type (**A**) and *prx9-1 prx40-2* (**B**) locules were compared by transmission electron microscopy. Tapetal cells are false colored in blue and developing pollen grains are false colored in orange. The *prx9-1 prx40-2* tapetum is swollen and hypervacuolated (black arrow). Pollen grains are also swollen and hypervacuolated (white arrow). Electron-dense sporopollenin-like material accumulates on the locular face of the tapetum (black arrowhead). Bars = 4 μm. Wild-type tapetal cells (**C**) have rectangular boundaries. *prx9-1 prx40-2* tapetal cells (**D**) have irregular, jigsaw-shaped boundaries (white arrowhead). Bars = 600 nm.

### PRX9 and PRX40 are active peroxidases *in vitro*

To probe the biochemical function of PRX9 and PRX40, we first tested whether PRX9 and PRX40 are catalytically active H_2_O_2_-dependent peroxidases by heterologously expressing PRX9 and PRX40 in *E. coli*. Although both proteins were highly expressed in *E. coli*, they were insoluble. We solubilized recombinant PRX9 and PRX40 by denaturing them in 8M urea. After purification, we subjected the unfolded PRX9 and PRX40 to a peroxidase refolding screen using pyrogallol as a generic peroxidase substrate (Smith et al., 1990). Refolding conditions were tested for activity by the addition of pyrogallol and H_2_O_2_. Activity was observed by a colorimetric change associated with an increase in absorbance at 420 nm as pyrogallol was converted into purpurogallin. We optimized the refolding conditions for both PRX9 and PRX40, and observed H_2_O_2_-dependent peroxidase activity on pyrogallol for both enzymes **(Figure. S3)**. This result demonstrates that PRX9 and PRX40 behave as typical class III peroxidases and are capable of completing the catalytic cycle in vitro.

### *PRX9* and *PRX40* are likely extensin cross-linking peroxidases

Since the cell wall exerts a strong influence on the size and shape of plant cells, and since class III peroxidases are typically localized in the plant apoplast, we hypothesized that PRX9 and PRX40 may oxidize a component of the tapetum and microspore cell walls and thus reinforce them. Since the tapetum is not thought to be lignified or suberized, extensins stood out as a potential substrate for PRX9 and PRX40. Extensins are structural hydroxyproline-rich glycoproteins in the cell wall that are known to regulate cell size and shape. They form large fibrillar networks and are thought to scaffold the assembly of the plant cell wall (Cannon et al., 2008). Previous studies showed that extensins may be intra- and inter-molecularly cross-linked by class III peroxidases via tyrosine residues (Brady et al., 1996; Brownleader et al., 1995; Fry, 1982). It is unclear why extensins must be cross-linked, but cross-linking is necessary for their function (Hall and Cannon, 2002; Ringli, 2010).

To assess the role of extensins in tapetal cell development, we first examined the accumulation of extensins in wild-type anthers by immunostaining. We took advantage of JIM20, a well-characterized extensin-specific monoclonal antibody (Pattathil et al., 2010). Consistent with our hypothesis JIM20 extensin epitopes were highly enriched in the wild-type tapetum (**Figure 6 A and B**). Pollen walls also appeared to be stained, but it is difficult to determine the extent of staining because pollen grains are naturally pigmented.

**Figure 6.**
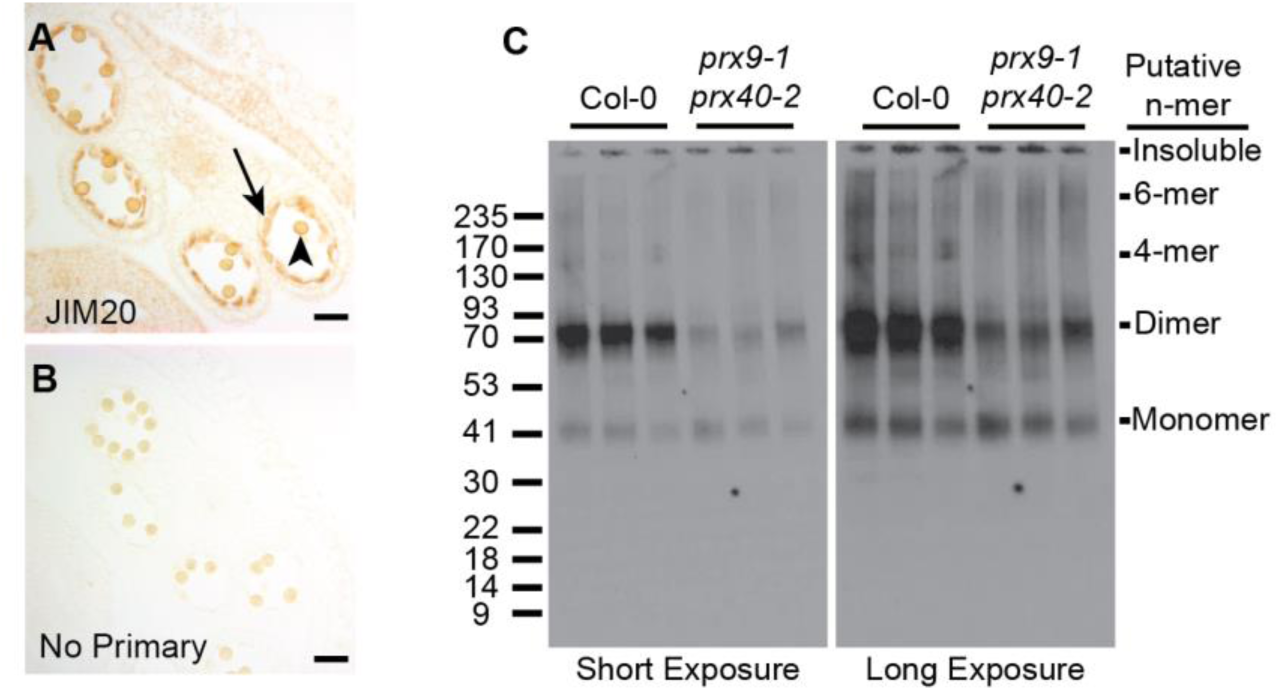
High-molecular-weight JIM20 extensin epitopes accumulate in wild-type but not *prx9-1 prx40-2* tapetal cells. Extensins in wild-type stage 10 anthers were examined by immunostaining with the JIM20 antibody (**A**), and compared to a control without a primary antibody (**B**). Staining was most obvious in the tapetum (black arrow) and may also be present in the pollen grain wall (black arrowhead). The apparent molecular weight of extensins were compared between three biological replicates of wild-type and *prx9-1 prx40-2* stage 10 - 12 (Sanders et al., 1999) anthers by western blot (**C**). Left panel is a short exposure and right panel is a long exposure of the same blot. Similar amounts of a putative ~40 kDa monomer are present in both samples. Significantly more putative ~80 kDa dimer are present in wild-type anthers. 160 kDa and 240 kDa species, corresponding to putative 4-mer and 6-mer, respectively, are present in wild-type anthers but absent from *prx9-1 prx40-2* anthers.

Since cross-linking increases the apparent molecular weight of extensins, we expected high-molecular-weight extensins to accumulate in the tapetum of wild-type anthers. If *PRX9* and *PRX40* were the primary extensin peroxidases in tapetum, then we would expect *prx9-1 prx40-2* anthers to accumulate less high-molecular-weight extensins. To test this hypothesis, we performed a western blot, comparing protein extracts of stage 10 (Sanders et al., 1999) wild-type and *prx9-1 prx40-2* anthers (**Figure 6C**). Wild-type anthers accumulate a putative monomeric extensin at approximately 40 kDa with higher-molecular-weight species also present. The most intense bands appeared to be at approximately 80 kDa, corresponding to the molecular weight of an extensin dimer. Additional bands at approximately 160 and 240 kDa were visible, likely corresponding to the molecular weight of a 4-mer and 6-mer, respectively. By comparison, *prx9-1 prx40-2* anthers accumulated significantly less 80 kDa extensins and the 160 kDa and 240 kDa bands were completely absent, even though *prx9-1 prx40-2* anthers accumulated approximately the same amount of the putative monomeric species. Altogether, these data provide *in vivo* evidences supporting the role of PRX9 and PRX40 as extensin cross-linking peroxidases in the tapetum.

### The *extensin18* mutant partially phenocopies *prx9 prx40* anthers

If extensin cross-linking by PRX9 and PRX40 is important for maintaining tapetum and microspore cell size, knock-out mutants of the extensins that are substrates for these peroxidases should phenocopy the *prx9 prx40* plants. Since no extensins in *Arabidopsis* were predicted to be specifically expressed in early stages of flower development, we could not identify specific extensin substrates for PRX9 and PRX40 by co-expression analysis. However, *EXTENSIN18* (*EXT18*) knock-outs have previously been reported to cause defects in male fertility (Choudhary et al., 2015). The *ext18* mutant was reported to exhibit decreased pollen viability, suggesting that EXT18 is likely expressed in developing anther and may be cross-linked by PRX9 and PRX40. To assess the role of *EXT18* in tapetum development, we performed a histological comparison between *ext18* (GT8324) and Ler-0 wild-type anther development (**Figure 7**). As previously reported, we observed degenerated pollen grains at later stages of anther development, but only very minor differences in the shape or size of the tapetum at earlier developmental stages. This result suggests that additional extensins likely serve as redundant or more prominent substrates for PRX9 and PRX40 in the tapetum.

**Figure 7.**
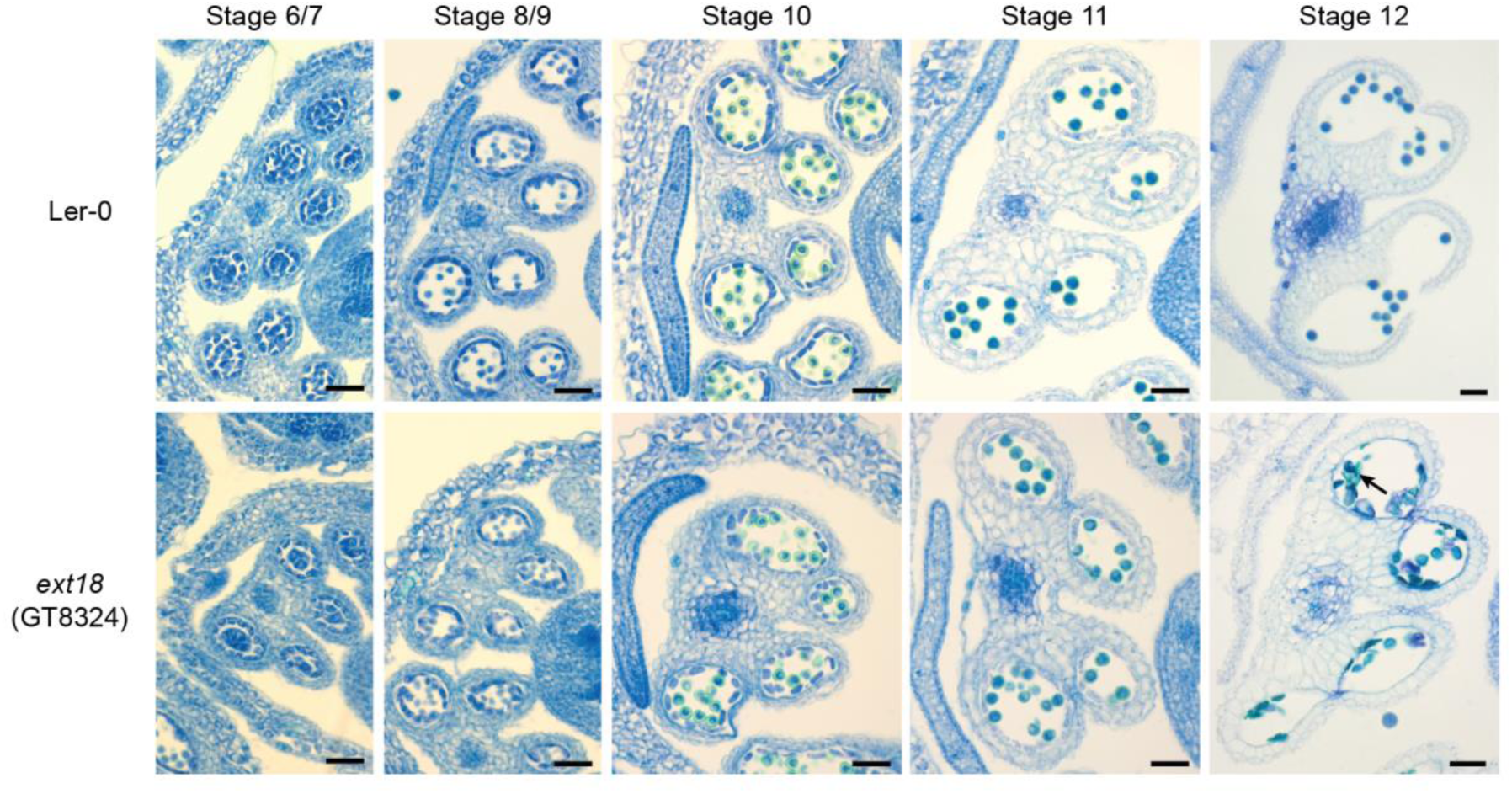
*ext18* anthers exhibit pollen degeneration but not tapetum hypertrophy. Wild-type (**Top Row**) and *ext18* (**Bottom Row**) anther development from stages 5 - 12 (Sanders et al., 1999). Wild-type and *ext18* are indistinguishable until stage 12 when numerous degenerated pollen grains are observed in *ext18* anthers (black arrow). Bars = 25 μm.

### PRX9 and PRX40 are evolutionarily conserved class III peroxidases in land plants

Since PRX9 and PRX40 play an essential role in *Arabidopsis* anther development, we sought to predict if orthologs are likely to be important in other plant species. To examine the evolution of PRX9 and PRX40, we performed comprehensive phylogenetic analyses of class III peroxidases across diverse land plant lineages. First, we performed a large-scale neighbor-joining phylogenetic analysis using 887 class III peroxidases collected from several reference model plants representing major land plant lineages (**Figure S4**). This initial analysis identified six major phylogenetic clades of peroxidases, termed clades i through vi, which overall agree well with previously published phylogenies (Tognolli et al., 2002). By mapping known functions associated with class III peroxidases to the tree, we observed that the phylogeny does not separate peroxidases by function. For example, lignin peroxidases do not cluster into one unique clade, but rather are distributed across clades i, iii, and vi. This may be a reflection of class III peroxidases’ well-recognized substrate promiscuity (Shigeto and Tsutsumi, 2016). Under low selective pressure to maintain substrate specificity, lignin peroxidase activities might have evolved multiple times in parallel during land plant evolution through gene duplication followed by neofunctionalization or subfunctionalization.

To inquire about the deep evolutionary history of PRX9 and PRX40, we then performed a more focused maximum-likelihood phylogenetic analysis using only the sequences from clade i, which contains both PRX9 and PRX40. This analysis revealed that *PRX40* is highly conserved among all major lineages of land plants, with orthologous sequences found in species that span bryophytes to angiosperms. In contrast, PRX9 falls into a smaller paralogous clade, which contains only angiosperm sequences. These results suggest that PRX40 orthologs likely emerged in the common ancestor of all land plants during early land colonization 450 million years ago, whereas *PRX9* emerged through parallel evolution much later within angiosperms.

## Discussion

### *PRX9* and *PRX40* function to maintain tapetum and microspore cell wall integrity during anther development

One of the most defining features of the *prx9 prx40* phenotype is tapetum hypertrophy and subsequent male-sterility. Tapetum hypertrophy is a defect in anther development that has been observed in many different contexts across multiple plant species. To our knowledge, tapetum hypertrophy was first reported in rice as a stress response to cold (Satake and Hayase, 1970; Mamun et al., 2006), and was later observed in cytoplasmic male-sterile *Capsicum annuum* and in heat- or abscisic acid-treated *Triticum aestivum* (Horner and Rogers, 1974; Saini et al., 1984). These early studies established a connection between environmental stress and tapetum hypertrophy-associated male sterility; however, the mechanistic basis for this phenomenon has remained elusive.

In the model plant *Arabidopsis thaliana*, tapetum hypertrophy has been seen as a developmental phenotype associated with a wide collection of mutants, including *roxy1*, *roxy2*, *ams, ms1, tdf1, dyt1, myb33, myb65, bhlh010, bhlh089, bhlh091, kns4/upex1*, *gne1, gne4*, and *fat tapetum* (Sorensen et al., 2003; Wilson et al., 2001; Zhu et al., 2008; Zhang et al., 2006; Sanders et al., 1999; Sorensen et al., 2002; Millar and Gubler, 2005; Suzuki et al., 2017; Phan et al., 2011; Xing and Zachgo, 2008; Zhu et al., 2015). Besides KNS4/UPEX1, which is a glycosyl transferase, these genes encode regulatory genes and transcription factors, or are unmapped mutant alleles. Hence, these mutants provided little insight towards a mechanistic understanding of tapetum hypertrophy or how it is prevented in the course of normal anther development in *Arabidopsis*. *PRX9* and *PRX40* are likely downstream of these transcriptional regulators and among the first effectors to be identified that directly influence tapetum size by cross-linking extensins in the cell wall.

### *PRX9* and *PRX40* are likely extensin peroxidases

Extensins are a large family of highly repetitive hydroxyproline-rich glycoproteins, which are ubiquitous structural cell wall proteins in land plants (Lamport et al., 2011). Extensins contain alternating hydrophilic and hydrophobic stretches that allow them to self-assemble into dendritic networks (Cannon et al., 2008). The characteristic hydrophilic extensin motif consists of heavily glycosylated Ser-(Hyp)_3-5_ interspersed with hydrophobic Tyr-Val-Tyr (Lamport et al., 2011). Class III peroxidases act on these tyrosines to form intra- and intermolecular cross-links of extensins.

The tapetal and microspore swelling phenotype seen in *prx9 prx40* anthers is reminiscent of other extensin-associated phenotypes. The most well-studied extensin in *Arabidopsis* is *EXT3/RSH (Hall and Cannon, 2002)*. Similar to *prx9 prx40* mutants, *rsh* mutants display defects in cell size and shape due to compromised cell walls. Cells within the developing embryos of *rsh* plants are disorganized and fail to form the characteristic heart shape (Hall and Cannon, 2002). Cells throughout seedling roots are also abnormally shaped and dysfunctional. Similarly, mutations in genes involved in extensin glycosylation display phenotypes associated with uninhibited cell expansion, including abnormally elongated roots and petioles, larger rosette leaves, and defects in root hair growth. (Gille et al., 2009; Møller et al., 2017; Velasquez et al., 2011; Saito et al., 2014).

Our experimental data also indicate that PRX9 and PRX40 play a role in regulating tapetum and microspore size and shape primarily by their extensin cross-linking activities. We observed the accumulation of extensins in the developing tapetum by immunostaining. Furthermore, we observed that *prx9 prx40* anthers failed to accumulate high-molecular-weight extensins, a direct consequence of extensin cross-linking *in vivo*. Finally, the pollen degeneration seen in *ext18* plants partially phenocopied the *prx9-1 prx40-1* phenotype.

Additionally, previous work has suggested that extensin disassembly is important for tapetal programmed cell death. A papain-like cysteine protease, *CEP1*, was previously found to be specifically expressed in the tapetum (Zhang et al., 2014). Papain-like cysteine proteases have the unique ability to degrade extensins during plant programmed cell death (PCD) (Helm et al., 2008). Indeed, the *Arabidopsis cep1* mutant displays delayed tapetal PCD, which may be partially explained by its inability to dismantle the extensin scaffold in the tapetum cell wall during late stages of anther development. Altogether, this evidence point to an important role for extensins in the tapetum and microspore cell walls.

### PRX9 and PRX40 substrates: Extensins and H_2_O_2_

Although EXT18 appears to be a substrate for PRX9 and PRX40, additional extensins are likely involved in the tapetum and microspore cell walls. Three extensin or extensin-like genes were found to be upregulated in *cep1* flower buds, implying their role in the tapetum: *EXT3/RSH/(AT1G21310), AT5G25550*, and *EXT21*/*AT2G43150 (Zhang et al., 2014)*. Of these three, only EXT3 and EXT21 contain the necessary tyrosines for cross-linking and are therefore possible candidate substrates for PRX9 and PRX40. In addition to *EXT18 (AT1G26250)*, *EXT19 (AT1G26240)* encodes another candidate extensin substrate. *EXT19* is a tandem duplicate of *EXT18* and the two share 84% sequence similarity at the protein level. *EXT18* and *EXT19* are arranged head-to-head in the genome, suggesting that they may be co-regulated by a bi-directional promoter. The roles for these extensins in tapetal and pollen walls are likely redundant or partially overlapping. Future work is necessary to dissect the functions of individual extensins and the coordination of these extensions during anther development.

As a co-substrate for class III peroxidases, H_2_O_2_ is required for the catalytic function of PRX9 and PRX40. H_2_O_2_ may be derived from multiple sources *in planta*. During plant defense and lignification, H_2_O_2_ is primarily derived from the reduction of O_2_ by the Respiratory Burst Oxidase Homologs (RBOHs) to superoxide (O_2_^−^) (Marino et al., 2012; Lee et al., 2013), which is further converted to H_2_O_2_ by superoxide dismutase (SOD). The *Arabidopsis* genome encodes 10 *RBOHs*: *RBOHA* through *RBOHJ. RBOHE* was shown to be specifically expressed in the tapetum during anther development (Xie et al., 2014). Interestingly, the *rbohe* mutant also displays a mildly hypertrophic tapetum, degenerated pollen grains, and pollen grains with distorted morphologies. Synergistic effects in *rbohe rbohc/rhd2* suggest that *RBOHC* may also play a role in generating tapetal H_2_O_2_. These phenotypes are consistent with *RBOHE* and *RBOHC* acting upstream of *PRX9* and *PRX40* by generating the O_2_^−^ which is subsequently converted to H_2_O_2_ by a putative SOD. At present, no tapetal *SOD* has been identified or characterized.

In summary, this work identifies *PRX9* and *PRX40* as critical genes for *Arabidopsis* anther development, likely through their extensin cross-linking activity. *PRX40* is highly conserved among all land plants, suggesting that extensin cross-linking was an ancient and critical function for the colonization of land by early land plants. *PRX9* likely arose later within angiosperms and descended from a duplication event that can be traced back to a *PRX40* progenitor. At present, most of the 73 class III peroxidases in *Arabidopsis* have no assigned functions. Future studies may identify additional class III peroxidases to be involved in extensin cross-linking during other developmental processes in plants. Finally, this work sheds light on the mechanism of tapetum hypertophy, a stress phenotype associated with reductions in crop yield of rice and wheat (Satake and Hayase, 1970; Mamun et al., 2006)(Horner and Rogers, 1974; Saini et al., 1984). As we continue to expand our understanding of the dynamic signaling and metabolic processes during anther development, we hope to improve our ability to develop stress-resistant crops.

## Materials and Methods

### Plant growth conditions

*Arabidopsis thaliana* Columbia (Col-0) seeds were placed on water-saturated 3:1 soil:vermiculite supplemented with Osmocote. Plants were grown in 16-hour light/8-hour dark days at 23 °C. T-DNA insertion Lines were obtained from the Arabidopsis Biological Resource Center or the Cold Spring Harbor Laboratory and grown under the same conditions.

### RT-PCR

Total RNA was extracted from approximately 100 mg of fresh plant tissue with the RNeasy mini kit (Qiagen) and subjected to an on-column DNase digest to remove contaminating genomic DNA. cDNA was prepared using the SuperScript™ III Reverse Transcriptase kit (Invitrogen) and used as template for RT-PCR.

### β-Glucuronidase Localization

The 507 bp *PRX9* promoter region and 959 bp *PRX40* promoter region were amplified out of Col-0 genomic DNA and ligated into BamHI-linearized pBI101.2 by Gibson Assembly. Assemblies were confirmed by DNA restriction digest and Sanger sequencing, and were transformed into *Agrobacterium tumefaciens* GV3101 strain. *Agrobacterium*-mediated *Arabidopsis* transformation was performed using the floral dip method (Clough and Bent, 1998). T1 kanamycin-resistant transformants were selected on Murashige and Skoog/Agar media containing 200 ug/mL timentin and 50 ug/mL kanamycin. After selection, kanamycin resistant seedlings were transferred to soil and grown under greenhouse conditions.

For GUS staining, flower buds from adult plants were collected into ice-cold 90% acetone and incubated for 20 minutes at room temperature. Samples were washed in staining buffer (50 mM Sodium phosphate pH 7.2, 0.2% Triton X-100, either 2 mM or 10 mM potassium ferrocyanide/potassium ferricyanide) and then incubated in staining solution (staining buffer containing 2 mM X-Gluc) on ice. Sample were infiltrated with staining solution under vacuum for 20 minutes, or until samples sank, and then incubated overnight at 37 °C to allow the GUS staining reaction to proceed. The following day, samples were processed through a graded series into 50% EtOH and placed in FAA (50% EtOH, 10% glacial acetic acid, 5% formaldehyde) for fixation. Finally, samples were placed in 70% EtOH for examination.

### Genotyping

Approximately 1 mg of plant tissue was placed in 50 uL of TE Buffer (10mM Tris pH 8, 1mM EDTA). Tissue was ground with a pestle and 1uL of the crude extract was used as a template for genotyping PCR. Genotyping primers were designed using the SIGnal Salk T-DNA primer design tool (http://signal.salk.edu/tdnaprimers.2.html).

### Paraffin Histology and Histochemistry

Formalin-Aceto-Alcohol (FAA)-fixed flower buds were transferred through a graded series into 100% EtOH and then through a grades series into 100% tert-butanol. Samples were then processed to paraplast and embedded in paraffin wax blocks. 8-10 micron sections were collected with a microtome, transferred to slides, and dried overnight at 37 °C. For GUS stained samples, slides were deparaffinized with two changes of Histo-clear and cover-slipped with permount. For Toluidine Blue staining, samples were deparaffinized with Histo-clear, rehydrated, stained with 0.5% Toluidine Blue, dehydrated and cover-slipped with permount.

### Scanning electron microscopy

Pollen grains were collected from open flowers and transferred to carbon adhesive tabs (Electron Microscopy Sciences) that were placed on aluminum mounts (Electron Microscopy sciences). Samples were sputter coated with Au/Pd with a Hummer sputter coater and imaged with Jeol 5600 LV SEM. For SEM of anthers, samples were fixed in 2% glutaraldehyde, 3% para-formaldehyde, 0.1M sodium cacodylate pH 7.4, 5% sucrose, 0.01% Triton X-100 overnight. Fixed samples were dehydrated through a graded series into 100% ethanol, critical point dried in a Tousimis critical point dryer, and sputter coated.

### Transmission electron microscopy

Stage 12 flower buds were fixed in 2.5% glutaraldehyde, 3% paraformaldehyde with 5% sucrose in 0.1M sodium cacodylate buffer (pH 7.4) and 0.01% Triton X-100 over-night. Samples were post fixed in 1% OsO_4_ in veronal-acetate buffer. Samples were stained *en block* overnight with 0.5% uranyl acetate in veronal-acetate buffer (pH6.0), dehydrated and embedded in Embed-812 resin. Sections were cut on a Leica EM UC7 ultramicrotome with a Diatome diamond knife at a thickness setting of 50 nm, stained with 2% uranyl acetate, and lead citrate. The sections were examined using a FEI Tecnai spirit at 80 KV and photographed with an AMT ccd camera.

### Immunostaining

10 micron sections were collected from paraffin embedded flower buds, deparaffinized with histoclear, and transferred into water through a graded series of ethanol. Sections were equilibrated in Dulbecco’s phosphate-buffered saline (PBS) and then blocked with 3% dry non-fat milk. 10 μL of undiluted JIM20 hybridoma supernatant (Carbosource) was added to the sections and incubated overnight at 4°C. Primary antibody was washed away with five exchanges of PBS. 10 μL of 1:10,000 mouse anti-Rat IgM secondary antibody-HRP (ThermoFisher) was added to sections and incubated at room temperature for one hour. Secondary antibody was washed away with five exchanges of PBS. 20 μL of 0.05% diaminobenzidine/0.015% H_2_O_2_ was added and allowed to incubate until sections showed robust staining. Reactions were stopped with a rapid dilution into milliQ and four subsequent exchanges with fresh milliQ.

### Western Blotting

Stages 10 - 12 anthers were dissected out of individual flower buds, frozen on dry ice, and ground in a microcentrifuge tube with a pestle. 50 μL of extraction buffer (25 mM Tris pH 8, 4% SDS) was added and the samples were mixed thoroughly with the pestle. Extracts were refrozen and stored at −20 °C. 10 μL samples were separated on Mops-Tris SDS-PAGE gels and transferred to nitrocellulose membranes. Membranes were blocked with 5% dry fat milk in TBS-T and incubated with 1:50 JIM20 overnight at 4 °C. Membranes were washed five times with TBS-T and incubated with 1:10,000 mouse anti-Rat IgM secondary antibody-HRP (Thermofisher). Immunostaining was visualized with the Enhance Chemiluminescence kit (Thermofisher).

### Recombinant Protein Expression and Refolding Optimization

*PRX9* and *PRX40* coding sequences were cloned without their predicted signal peptide sequences peptides into the pHIS8-4 bacterial expression vector with an N-terminal His-tag and transformed into *E. coli* BL-21(DE3). 500 mL of transformed bacteria were grown in Terrific Broth at 37°C/225 RPM until reaching OD_600_ of 0.8. Cultures were induced with 500 μM IPTG and incubated overnight at 16 °C/225 RPM. Cultures were pelleted by centrifugation and pellets were resuspended in 12 mL of lysis buffer (50 mM Tris pH 8, 500 mM NaCl, 30 mM Imidazole). Resuspensions were divided into 1 mL aliquots, frozen, and stored at −80 °C.

Resuspensions were thawed and lysed by sonication for 20 seconds. Lysates were centrifuged for 30 minutes at 16,000 RPM and supernatants were discarded. Insoluble PRX9 and PRX40 were washed 3 times by resuspending the pellets in lysis buffer with 2 M Urea, spinning for 30 minutes at 16,000 RPM, and discarding the supernatant. PRX9 and PRX40 were then solubilized in lysis buffer with 8M Urea and bound to Ni^2+^-NTA resin to separate them from bacterial cell material. Protein-bound Ni^2+^-NTA resin was diluted into elution buffer (50 mM Tris pH 8, 500 mM NaCl, 300 mM Imidazole) with 8M Urea and the resin was separated from the elution by centrifugation.

Refolding conditions were tested by diluting 10 μL of of His-Trap elution into 190 μL of various refolding conditions in a 96-well plate format. Conditions were sealed with an adhesive plate cover and refolding conditions were allowed to incubate for 60 hours in the dark at room temperature. 50 μL of refolding reactions were added to 50 μL of pyrogallol reaction buffer (final concentration: 10 mM Mes pH 6, 50 mM pyrogallol, 1 mM H_2_O_2_). The relative activity of refolding conditions was assessed by monitoring the formation of purpurogallin, which is yellow and absorbs light at 420 nm.

### Phylogenetic Analysis

Protein sequences from *Arabidopsis thaliana*, *Medicago truncatula*, *Oryza sativa, Zea mays, Selaginella Moellendorffii, Physcomitrella patens*, and *Marchantia polymorpha* genomes, were collected from Phytozome (https://phytozome.jgi.doe.gov). *Picea abies* sequences were collected from Congenie (http://congenie.org/). Sequences were aligned in MEGA6 (Tamura et al., 16AD) with the MUSCLE algorithm (Edgar, 2004). For the large-scale phylogeny **(Figure S4)** all 877 sequences were included. The tree was generated with the neighbor-joining method (Saitou and Nei, 1987). Evolutionary distances were calculated using the JTT matrix-based method (Jones et al., 1992) and are in units of number of amino acid substitutions per site.

For the maximum likelihood tree **(Figure 8)**, only the sequences from clade i of the neighbor-joining tree were included. Sequences were realigned in MEGA6 with MUSCLE. Positions in the alignment that were represented by only a single sequence were manually removed and the truncated sequences were realigned. A distance matrix was generated in MEGA6 and sequences with high divergence from *PRX9* and *PRX40* were manually removed. Several preliminary phylogenies were generated and sequences that could not be confidently placed on branches that related to *PRX9* or *PRX40* were manually removed. The final phylogeny was generated with the remaining sequences using the maximum likelihood method based on the Whelan And Goldman (WAG) model (Whelan and Goldman, 2001). A discrete Gamma distribution was used to model evolutionary rate differences among sites (5 categories (+G, parameter = 0.7211)). Branch lengths were measured in the number of substitutions per site.

**Figure 8.**
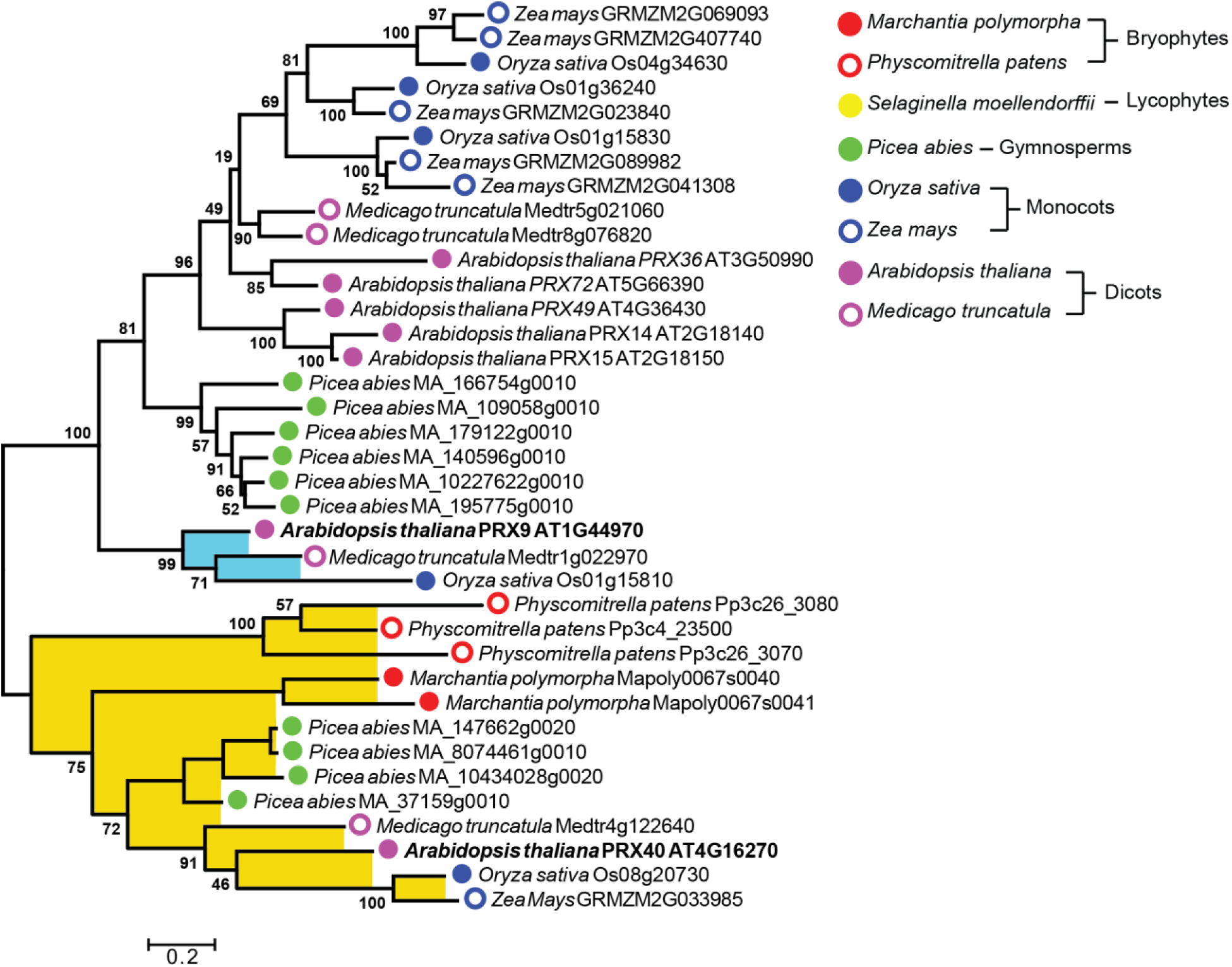
Maximum-likelihood phylogenetic analysis of *PRX9*, *PRX40*, and related class III peroxidases from various plant lineages. The analysis was generated with 1000 bootstrap replications using a subset of sequences from clade i (**Figure S4**). Bootstrap values are indicated at nodes within the tree. The scale measures evolutionary distance in substitutions per amino acid. *PRX9* is placed within a clade containing only angiosperm sequences, highlighted in blue. *PRX40* is placed in a clade containing sequences spanning land plants from angiosperms to bryophytes, highlighted in yellow. *PRX9* and *PRX40* branch labels are bolded.

## Accession Numbers

Sequence data from this article can be found in the EMBL/GenBank data libraries under accession numbers AT1G44970 (*PRX9*), AT4G16270 (*PRX40*), AT3G18780 (*ACTIN2*) AT1G26250 (*EXT18*)

## Supplemental Figures

**Supplemental Figure 1.** BAR eFP predicts *PRX9* and *PRX40* are expressed in early flower buds.

**Supplemental Figure 2.** Anther filaments from stage 14 *prx9-1 prx40-2* flower are not fully elongated.

**Supplemental Figure 3.** Reconstituted enzyme activity can be obtained by solubilizing, purifying, and refolding protein from *E. coli* expressing PRX9 and PRX40.

**Supplemental Figure 4.** Neighbor-joining phylogenetic analysis of class III peroxidases across land plants.

**Supplemental Table 1.** List of primers used in this study.

**Supplemental File 1.** Alignment of class III peroxidases protein sequences used for large-scale neighbor-joining phylogenetic analysis **(Figure S4)**.

**Supplemental File 2.** Alignment of sequences used for maximum-likelihood analysis **(Figure 8)**.

## Acknowledgements

We thank Nicki Watson at the Whitehead Institute Keck Imaging Facility for assistance with electron microscopy. This work was supported by the Pew Scholars Program in the Biomedical Sciences (grant number 27345) and the Searle Scholars Program (grant number 15-SSP-162). This material is also based upon work supported by the National Science Foundation Graduate Research Fellowship under Grant No. 1122374.

## Author Contributions

J.R.J. and J.K.W. designed and analyzed all experiments. J.R.J. performed the experiments. J.R.J. and J.K.W. wrote the paper.

